# Homology-directed repair of a defective *glabrous* gene in *Arabidopsis* with Cas9-based gene targeting

**DOI:** 10.1101/243675

**Authors:** Florian Hahn, Marion Eisenhut, Otho Mantegazza, Andreas P.M. Weber

## Abstract

The CRISPR/Cas9 system has emerged as a powerful tool for targeted genome editing in plants and beyond. Double-strand breaks induced by the Cas9 enzyme are repaired by the cell’s own repair machinery either by the non-homologous end joining pathway or by homologous recombination. While the first repair mechanism results in random mutations at the double-strand break site, homologous recombination uses the genetic information from a highly homologous repair template as blueprint for repair of the break. By offering an artificial repair template, this pathway can be exploited to introduce specific changes at a site of choice in the genome. However, frequencies of double-strand break repair by homologous recombination are very low. In this study, we compared two methods that have been reported to enhance frequencies of homologous recombination in plants. The first method boosts the repair template availability through the formation of viral replicons, the second method makes use of an *in planta* gene targeting approach. Additionally, we comparatively applied a nickase instead of a nuclease for target strand priming. To allow easy, visual detection of homologous recombination events, we aimed at restoring trichome formation in a glabrous *Arabidopsis* mutant by repairing a defective *glabrous1* gene. Using this efficient visual marker, we were able to regenerate plants repaired by homologous recombination at frequencies of 0.12% using the *in planta* gene targeting approach, while both approaches using viral replicons did not yield any trichome-bearing plants.

## INTRODUCTION

Gene targeting (GT) means integration of foreign DNA into a cell’s genome by homologous recombination (HR; Paszkowski et al., 1988). GT has been a long-term goal for plant scientists since it allows modifying an endogenous gene *in planta* or integrating a transgene at a specific position in the genome (Puchta, 2002). GT is easily achieved in lower eukaryotes, such as yeast, or the moss *Physcomitrella patens,* because they include foreign DNA predominantly through a highly efficient HR pathways. However, GT is not easily achieved in higher plants, since they integrate foreign DNA via illegitimate recombination through the non-homologous end joining (NHEJ) pathway (Puchta, 2005). A simple introduction of a DNA donor molecule with homology arms (HA) only led to frequencies of 10^−4^ to 10^−6^ GT events per transformation event in higher plants, such as tobacco and *Arabidopsis* (Lee et al., 1990; Offringa et al., 1990; Miao and Lam, 1995; Risseeuw et al., 1995), making GT impractical for most applications. Nevertheless, GT frequencies in higher plants can be increased by two orders of magnitude by inducing double-strand breaks (DSB) in the target locus with sequence specific nucleases (SSN, Puchta et al., 1996). SSN are proteins that can cleave DNA at predefined sequence motives. The first generation of SSN, including meganucleases, Zinc-finger nucleases (ZFN), and Transcription Activator-like Effector Nucleases (TALENs) recognized DNA by protein-DNA interactions, which required rational protein design for every target sequence (Voytas, 2013). The most recent addition to the SSN has been the CRISPR/Cas9 system (Cas9 system).

Cas9 is an endonuclease of the bacterium *Streptococcus pyogenes* that generates DSB in DNA (Doudna and Charpentier, 2014) and that recently has become the SSN of choice for genome editing and GT. Cas9 became so popular because, in contrast to other SSNs that recognize DNA sequences through protein-DNA interactions, Cas9 is guided to its target site on DNA by complementarity matching of a single guide RNA (sgRNA, Jinek et al., 2012). The RNA-based targeting system makes Cas9 easily programmable towards a selected target. Therefore, the Cas9 system has been used extensively by numerous laboratories to edit plant genomes (Zhang et al., 2017). Additionally, mutation of one of the two endonuclease sites of Cas9 resulted into a Cas9 nickase, which introduces only single-strand nicks in the plant genome (Fauser et al., 2014).

SSN-generated DSB are lethal for the cell and must be repaired by dedicated protein machineries. Two major repair pathways can be distinguished: NHEJ and HR (Puchta, 2005). Most DSB in higher eukaryotes are repaired by NHEJ (Sargent et al., 1997), a pathway that involves direct re-ligation of the DSB ends (Gorbunova and Levy, 1997; Dueva and Iliakis, 2013), making it an error-prone process. This process can be exploited to introduce random small indel mutations at a DSB site and has been used extensively to knock-out gene functions with SSN (Bortesi and Fischer, 2015). In contrast, HR repair pathways use a homology template (HT) to repair a DSB and are therefore of high potential for GT. In the conservative synthesis-dependent strand annealing pathway, which is the most prominent HR pathway in plant somatic tissues (Puchta, 1998), DNA ends are resected to create 3’ overhangs. One 3’-end can invade into a homologous sequence forming a loop-structure and copy the sequence from the HT DNA. After release from the loop structure, the single strand can reanneal with homologous sequences on the opposite side of the DSB, leading to conservative repair without sequence information loss (Steinert et al., 2016). Artificial HT with HA have already been used for GT experiments to integrate foreign DNA sequences into a DSB generated by SSN (Puchta et al., 1996; Baltes et al., 2014; Schiml et al., 2014; Li et al., 2015).

While GT frequencies were successfully enhanced by cutting the target site with SSN, this is not sufficient for streamlined application of GT in plants. GT frequencies can be further enhanced of approximately two orders of magnitudes (compared to conventional T-DNA delivery) by mobilizing the HT after T-DNA transformation. This can be achieved with an *in planta* gene targeting (IPGT) approach and with a viral replicon approach.

The IPGT approach (Fauser et al., 2012) employs SSN not only to cut the target locus but also to excise the HT from the chromosomal T-DNA backbone. The liberation from the genomic context of the T-DNA allows the activated HT to be available at the site of the DSB where it can function as repair template. This method has been used successfully using various SSN in *Arabidopsis* (Fauser et al., 2012; Schiml et al., 2014; Zhao et al., 2016), maize (Ayar et al., 2013; Kumar et al., 2016), and rice (Sun et al., 2016).

An alternative approach has been developed by Baltes et al. in tobacco. It activates the HT by including it in a *Geminivirus* based replicon (Baltes et al., 2014). Therefore, the HT is flanked with three viral elements from the *Bean yellow dwarf virus* (BeYDV) that are required to establish the replicon. These regions are known as the cis-acting long intergenic region (LIR), the short interacting region (SIR) and the gene for the trans-acting viral replicase *Rep/RepA.* The Rep protein initiates a rolling circle endoreplication at the LIR site, leading to a replicational release of the HT in form of up to several thousand circular replicons (Chen et al., 2011). This method has been successfully applied using components from various virus strains and with different SSN for GT in wheat protoplast (Gil-Humanes et al., 2017), tomato (Čermák et al., 2015), potato (Butler et al., 2016), and rice (Wang et al., 2017). While IPGT has already been shown to induce stable GT events in *Arabidopsis* (Fauser et al., 2012; Schiml et al., 2014), reports about successful applications of viral replicons based GT approaches in *Arabidopsis* have not been published so far.

In this report, we aimed to evaluate and compare the efficiencies of the viral replicon system and the IPGT system to enhance GT frequencies in the model plant *Arabidopsis thaliana.* Therefore, we made use of the R2R3-MYB transcriptional master regulator of trichome formation GLABROUS1 (GL1), which we established previously as non-invasive, endogenous visual marker for Cas9 activity in *Arabidopsis* (Hahn et al., 2017). Null-mutations of the *GL1* gene lead to loss of trichome formation (Koornneef et al., 1982) on stems and leaves as GL1 is part of the signaling cascade which allows epidermal cells to differentiate into trichomes (Pattanaik et al., 2014). Using the IPGT approach and the viral replicon-based approach, we aimed at repairing a non-functional *gl1* gene with a 10 bp deletion in the coding region (Hahn et al., 2017) by HR and by this, restoring trichome formation. Using IPGT, we achieved successful GT and regenerated three stable non-glabrous plants out of ~2,500 plants screened over three generations. However, we were not able to regenerate non-glabrous plants using viral replicons. The *gl1* phenotype provided a simple and efficient visual marker for tracking mutagenic activity *in planta.*

## METHODS

### Plant growth conditions

Surface-sterilized seeds of an *Arabidopsis gl1* mutant with a 10 bp deletion in the *GL1* gene (Hahn et al., 2017) were stratified at 4 °C for three days. Seeds were germinated on 0.8% (w/v) agar-solidified half-strength Murashige and Skoog medium (Duchefa) in growth chambers with a light intensity of 100 μmol photons m^−2^ s^−1^ under long day conditions (16-h-light/8-h-dark cycle, 22/18 °C). After 14 days, seedlings were transferred to soil and further grown under the same conditions. For large-scale screening of plants in the T2 generation, seeds were directly sowed on soil and the plants were grown in the green house after three days of stratification at 4 °C.

### Generation of a homology repair template for reparation of the ten nucleotides deletion in the *gl1* gene

We designed a homology repair template (HT) containing a codon-modified version of the missing 10 bp. The 10 bp (CTGCCGTTTA) were flanked by homology arms (HA) complementary to the genomic regions adjacent to the cutting site of Cas9 in the *gl1* gene. The HA consisted of 813 bp and 830 bp, respectively. Both homology arms contained a single nucleotide exchange within the homology region to distinguish two-sided true GT events from one-sided GT events combined with NHEJ mediated repair pathways (Schiml et al., 2014). After a short linker sequence, the sequence of the sgRNA recognition site including the protospacer adjacent motif (PAM) sequence followed. The HT was synthesized at GeneART (ThermoFisher Scientific) and integrated in the vector backbone pMA-RQ (GeneART). The resulting vector was named pFH95.

### Cloning of a binary T-DNA repair vector for *in planta* gene targeting repair approach

We made use of the binary T-DNA vector pDE-Cas9, containing an *Arabidopsis*-codon-optimized Cas9 controlled by the constitutive *Ubiquitin* promoter (PcUBp) from *Petroselinum crispum* (Fauser et al., 2014) and the sgRNA subcloning vector pEN-C.1.1 containing the *Arabidopsis* U6–26 promoter for sgRNA expression, the sgRNA scaffold and BbsI-sites for subcloning of the protospacer sequence (Schiml et al., 2014). After digestion of pEN-C1.1 with BbsI-HF (NewEngland Biolabs), we inserted the target sequence (AAGAGATGTGGGAAAAGAGA plus an additional guanine as first bp for transcription by the U6–26 promoter) using two annealed primers FH210/FH211 (Table S1) resulting in vector pFH89. We then digested pFH89 with MluI-HF (NEB) to extract the sgRNA cassette and ligated it into MluI-HF-digested pDE-Cas9, thereby creating the vector pFH99. We amplified the HT from pFH95 using primers FH278/FH279 (Table S1) and performed Gateway^®^ BP-reaction with pDONR 207 (Invitrogen) resulting in vector pFH122. LR reaction (Invitrogen) of pFH99 with pFH122 led to the assembly of the IPGT repair vector pIPGT-Nuc.

### Generation of binary T-DNA repair vectors for viral replicon repair approach

For the viral replicon repair approach, we amplified the viral components from BeYDV necessary for replicon formation from the vector pZLSLD.R (Baltes et al., 2014) using primers FH217/FH218 (Table S1) and cloned the PCR product into pDONR 207 (Invitrogen) via BP reaction. This vector (pFH83) was then digested with PmII (NewEngland Biolabs) and BsaXI (NewEngland Biolabs) to remove the original HT from Baltes et al. (2014). We PCR-amplified the *GL1* HT from pFH95 using primers FH240/FH241 (Table S1) and inserted it into the digested pFH83 vector by Gibson^®^ cloning (NewEngland Biolabs), resulting in the vector pFH84. The final viral replicon nuclease repair construct pVIR-Nuc containing Cas9 nuclease, the sgRNA cassette directed against the *gl1* gene and the HT was then cloned by Gateway^®^ LR reaction (Invitrogen) of pFH99 with pFH84.

For the viral replicon repair approach using the Cas9 nickase, we used the MluI-HF-extracted sgRNA cassette from pFH89 and ligated it into MluI-HF-digested pDE-Cas9-D10A (Fauser et al., 2014) resulting in the vector pFH103. LR reaction of pFH103 with pFH84 led to the final viral replicon nickase repair construct pVIR-Nick.

The sequences of the vectors pIPGT-Nuc (accession number xxx), pVIR-Nuc (accession number xxx) and pVIR-Nick (accession number xxx) are accessible on GenBank.

### Plant transformation

*Agrobacterium tumefaciens,* strain GV3101::pMP90 was transformed with the vectors pIPGT-Nuc, pVIR-Nuc, and pVIR-Nick. Floral dipping (Clough and Bent, 1998) was used to transform *Arabidopsis gl1* plants with the three constructs. Kanamycin (pFH136) and glufosinate (pFH114, pFH135) were used for selection of primary transformants. Furthermore, leaf genomic DNA was isolated from surviving plants via isopropanol precipitation (Weigel and Glazebrook, 2009). The presence of T-DNA insertion was verified by PCR using primer pairs FH190/FH223 (pIPGT-Nuc) or FH223/FH265 (pVIR-Nuc, pVIR-Nick, Table S1).

### Detection of the *Cas9* gene in plants

Presence of the *Cas9* gene in plant genomes was verified by PCR on leaf genomic DNA using primer pairs amplifying specifically the *Cas9* gene (FH313/FH314, Table S1).

### Verification of viral replicon formation

Formation of circular viral replicons was verified in T1 generation plants by PCR using leaf genomic DNA and the primers FH266/FH271 (Table S1).

### Detection of Cas9-mediated gene repair

For verification of HDR events on genomic level, leaf genomic DNA was isolated via isopropanol precipitation (Weigel and Glazebrook, 2009) and the region of interest in the *gl1* gene was PCR-amplified with proofreading Q5© polymerase (NewEngland Biolabs) using primers FH214/FH215 (Table S1). The obtained PCR amplicons were analyzed directly by Sanger sequencing (Macrogen) or subcloned into pJET1.2 (ThermoFisher Scientific) and then sequenced. 4Peaks software (Nucleobytes) and SnapGene Viewer (GSL Biotech) were used to analyze sequencing histograms.

## RESULTS

### Selection of a visual marker for testing gene targeting events

In a previous study, we established trichome formation as efficient visual marker to monitor Cas9 mutagenic activity in *Arabidopsis* (Hahn et al., 2017). We used a CRISPR/Cas9 approach to generate a T-DNA free glabrous *Arabidopsis* mutant that contains a deletion of 10 bp in the master regulator gene for trichome formation, *GL1.* This 10 bp deletion disrupts the reading frame of *GL1* and leads to a premature stop codon. Homozygous mutant plants show almost no leaf and stem trichomes (Fig. S1). We decided to use this mutant line as a model system to test Cas9-based GT in *Arabidopsis.* Cas9-induced cleavage in combination with homology directed repair can be used to integrate sequences of choice into a genomic region. If we successfully reintegrated the missing 10 bp into this mutant line’s genome, we would expect restoration of leaf trichome formation. Since the mutant line was created by CRISPR/Cas9, a PAM sequence was already present at the site of mutation and the predicted Cas9 cleavage site (3 bp upstream of the PAM sequence) matched the desired integration site for the missing 10 bp (Fig. S1).

### Selection of a gene targeting system and design of related constructs

HR is more efficient in plants if the HT is available at the site of the DSB (Baltes et al., 2014). If the HT is delivered as T-DNA, which integrates randomly into the genome, it might not be available at the DSB site for the DNA repair machinery. Thus, we comparatively analyzed two methods to enhance the availability of the HT at the DSB site.

The first method was developed by Baltes et al (2014) in tobacco calli. This method increases the availability of the HT by including it in small circular geminiviral replicons.

For the repair of the *gl1* gene, we accordingly created a single T-DNA vector, (Fig. 1A), that contained an *Arabidopsis* codon-optimized *Cas9* gene under the control of the constitutive PcUB promoter (Fauser et al., 2014), a sgRNA cassette targeting the *gl1* gene, and the HT with ~ 800 bp of HA on both sides, flanked by LIRs, SIR, and a *Replicase A* (*RepA*) gene. For an easy nomenclature of the plasmids, we decided to include the name of the HT activation method (viral replicons = vir; *in planta* gene targeting = IPGT) and the used Cas9 variant (nuclease = Nuc; nickase = Nick) in the plasmid name and therefore named this construct pVIR-Nuc.

**Figure 1:**
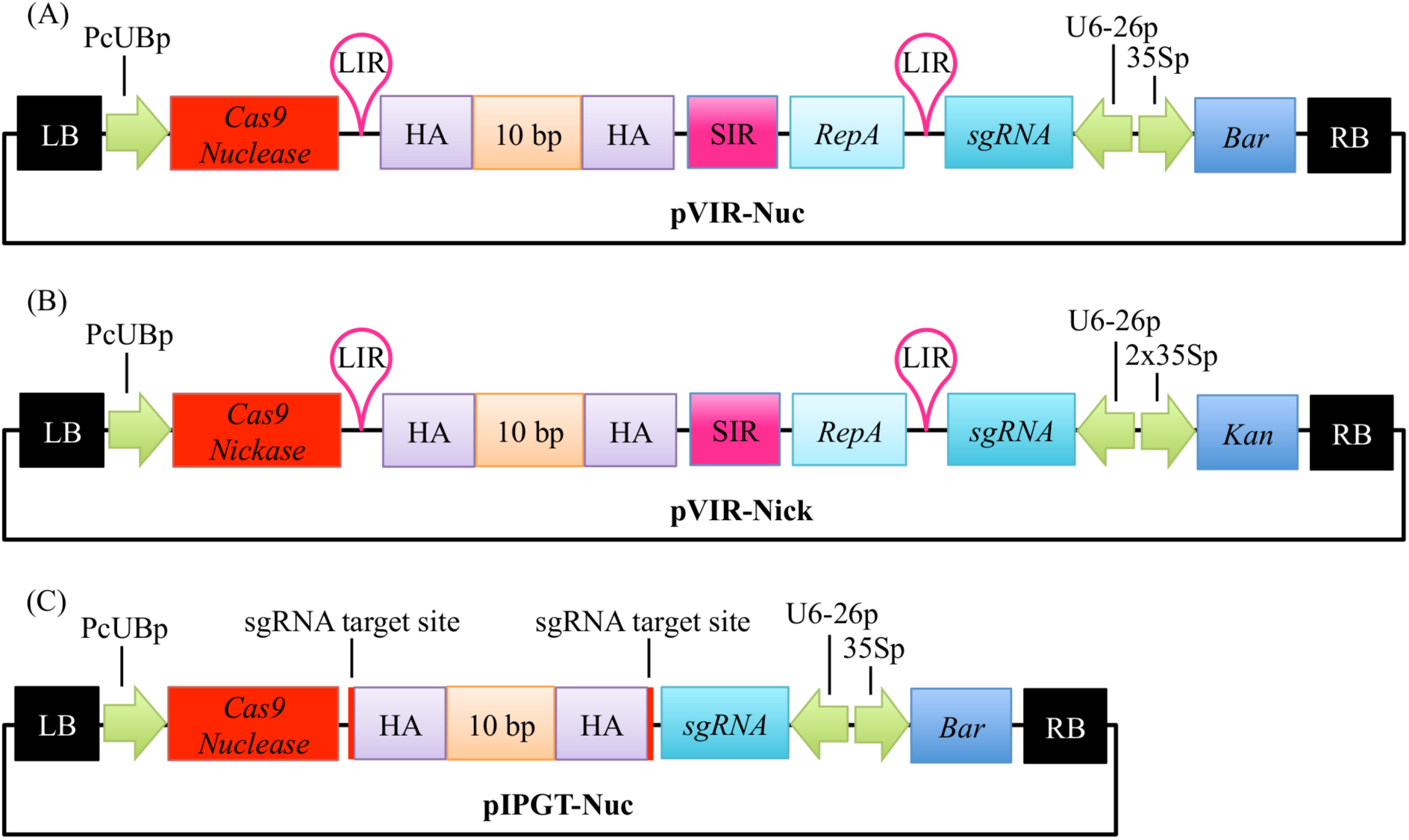
Vector design for repair of the *gl1* gene. Single T-DNA vectors were designed to repair the dysfunctional *gl1* gene. (A) pVIR-Nuc contains the Cas9 Nuclease gene (red box) under a constitutive *Ubiquitin6* promoter from parsley (PcUBp) and the sgRNA cassette (petrol box) controlled by the *U6–26* promoter (U6–26p). A Basta resistance cassette (*Bar,* dark blue box) is included as selection marker. The repair template itself consists of 10 nucleotides (10 bp, CTGCCGTTTA, orange box), which should restore the reading frame of the *gl1* gene flanked by homology arms (HA, purple box). Rolling circle replication of the HT is ensured by the flanking long intergenic region (LIR, pink hairpin), short intergenic region (pink box, SIR), and the gene encoding the replicase *RepA* (light blue box). (B) pVIR-Nick contains similar features but harbors the gene for the Cas9 nickase instead of the nuclease and a kanamycin resistance cassette (*Kan*, dark blue box) as selection marker. (C) pIPGT-Nuc contains the gene for the Cas9 nuclease, the sgRNA cassette, the Basta resistance cassette and the repair template with HA flanked by target sites for the Cas9 nuclease (red lines). RB = right T-DNA border, 35Sp = *Cauliflower Mosaic Virus* 35S promoter, LB = left T-DNA border. Size not to scale.

After expression, RepA binds to the LIR sites and initiates rolling circle replication, leading to the formation of up to several thousand copies of the HT in the nucleus (Timmermans et al., 1992; Baltes et al., 2014), which can then be used as repair template for the DSB created by Cas9 nuclease (Fig. 2A).

**Figure 2:**
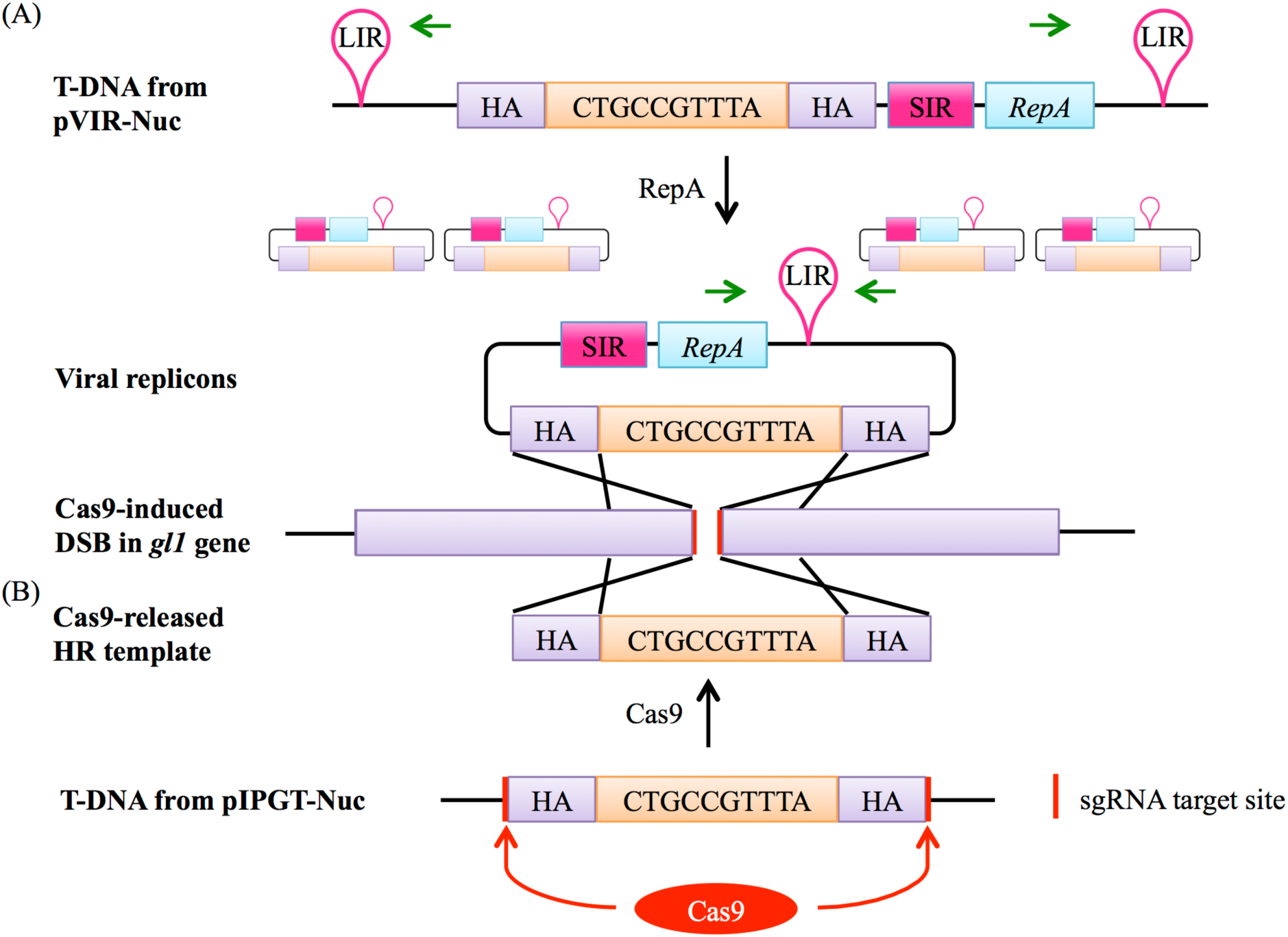
Two different mechanisms enhance the availability of the HT in the plant cell nucleus. (A) If the T-DNA from pVIR-Nuc is integrated into the plant genome, RepA is expressed and initiates rolling circle endoreplication of the HT leading to thousands of viral replicons. The presence of viral replicons can be detected by PCR using primers that only generate an amplicon on the circular replicons but not on the T-DNA (green arrows). The HT can then attach to the DSB site in the *gl1* gene induced by Cas9 due to homology between the HA and the genomic regions next to the DSB (purple). Repair of the DSB by HR will then lead to integration of the 10 bp, which restores the ORF. pVIR-Nick functions accordingly, except that Cas9 nickase only induces a DNA nick in the *gl1* gene. (B) The HT in the T-DNA of pIPGT-Nuc is flanked by the same sgRNA target site that is present in the *gl1* gene. Expression of Cas9 will therefore simultaneously target the *gl1* gene and release the HT, which can then attach to the DSB and function as repair template.

To further explore the potential of this method, we additionally generated and tested a complementary vector pVIR-Nick in which the Cas9 nuclease gene was replaced by a nickase gene (Fig. 1B). DNA nicks trigger HR without triggering the competing NHEJ repair (Fauser et al., 2014). NHEJ repair interferes with HR because it induces random indel mutations that destroy the sgRNA target site and make it unsuitable for homology directed repair. By using the Cas9 nickase, we expected to increase HR efficiency.

The second method, developed by Fauser et al. (2012), is called IPGT. This method increases the availability of the HT by releasing it from the T-DNA backbone using the same SSN that creates the DSB in the target region. Therefore, we designed the vector pIPGT-Nuc (Fig. 1C), in which the HT is flanked by the same sgRNA target site that is used to create the DSB in the *gl1* gene (Fig. 2B).

### Plants transformed with pVIR-constructs form circular viral replicons

We transformed Arabidopsis *gl1* plants with either pVIR-Nuc, pVIR-Nick, or pIPGT-Nuc and selected three independent T1 lines per construct. Though we did not detect trichomes on the leaf surface of any T1 plant, we wanted to verify that the viral components were indeed producing circular HT in the T1 plants. Thus, we performed a PCR using primers that only yield a product in the circularized version of the HT but not in the linear T-DNA (Fig. 2A). We detected PCR products in the plants transformed with pVIR-Nuc and pVIR-Nick. PCR fragments for plants transformed with pIPGT-Nuc, which do not generate replicons (Fig. S2), were not obtained.

### T2 generation plants show spots of trichomes on leaves

We analyzed several hundred plants from the progeny of each T1 plant for trichome reappearance. While we could not detect plants that fully recovered trichome production on leaves, we found several plants from the IPGT approach that harbored spots of trichomes (Tab. 1). These spots ranged from two trichomes on a single leaf of the plant to large spots on several leaves (Fig. 3A). In contrast, using the viral replicon method, we only detected one plant with a single trichome in plants transformed with pVIR-Nuc, which implements a full Cas9 nuclease to trigger HR. Plants transformed with pVIR-Nick (Fig. 3A), which implements the Cas9 nickase, did not form any trichomes on their leaves.

**Table 1:**
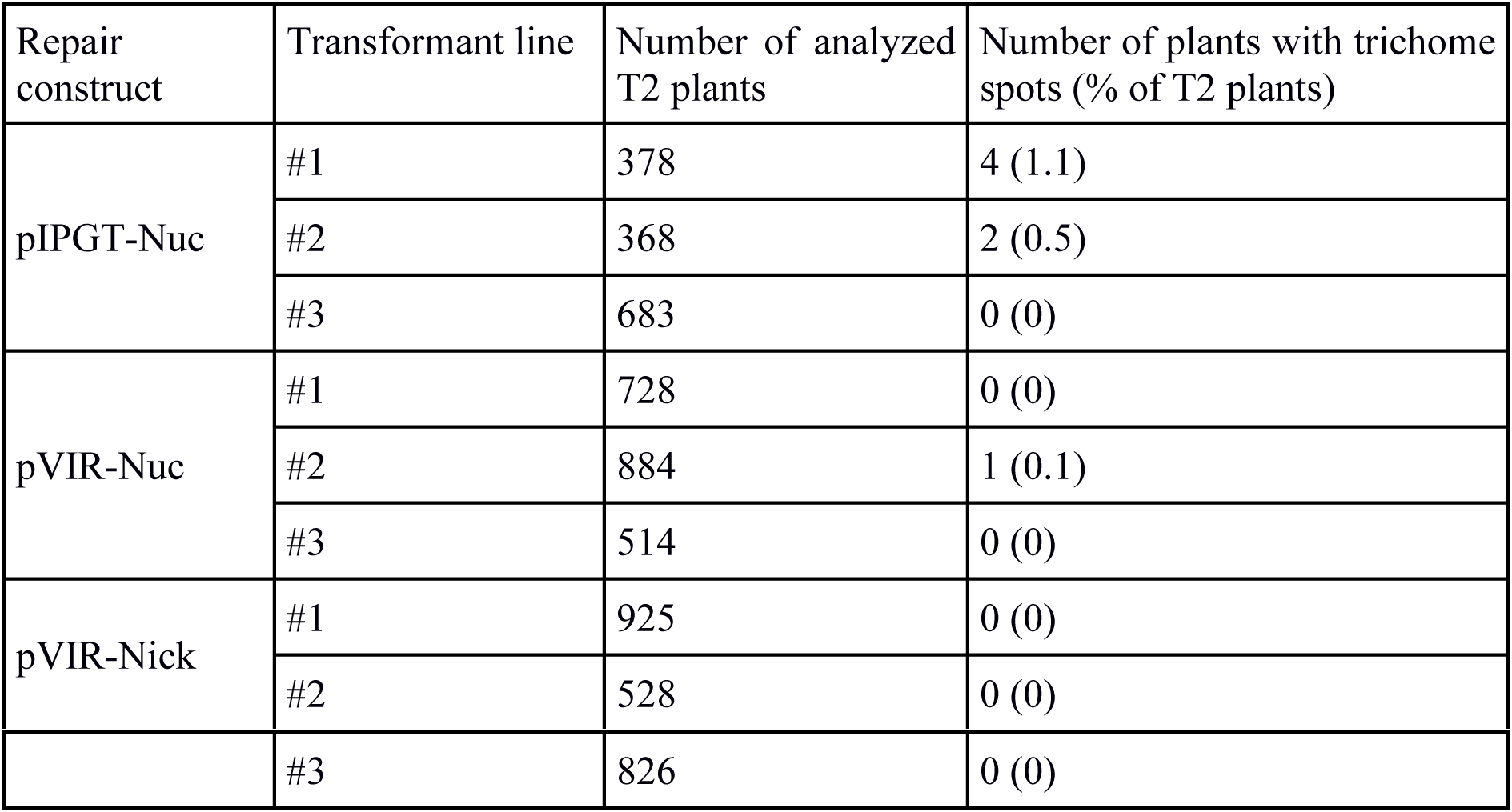
Trichome appearance in the T2 generation.

| Repair construct | Transformant line | Number of analyzed T2 plants | Number of plants with trichome spots (% of T2 plants) |
| --- | --- | --- | --- |
| pIPGT-Nuc | #1 | 378 | 4 (1.1) |
|  | #2 | 368 | 2 (0.5) |
|  | #3 | 683 | 0 (0) |
| pVIR-Nuc | #1 | 728 | 0 (0) |
|  | #2 | 884 | 1 (0.1) |
|  | #3 | 514 | 0 (0) |
| pVIR-Nick | #1 | 925 | 0 (0) |
|  | #2 | 528 | 0 (0) |
|  | #3 | 826 | 0 (0) |

**Figure 3:**
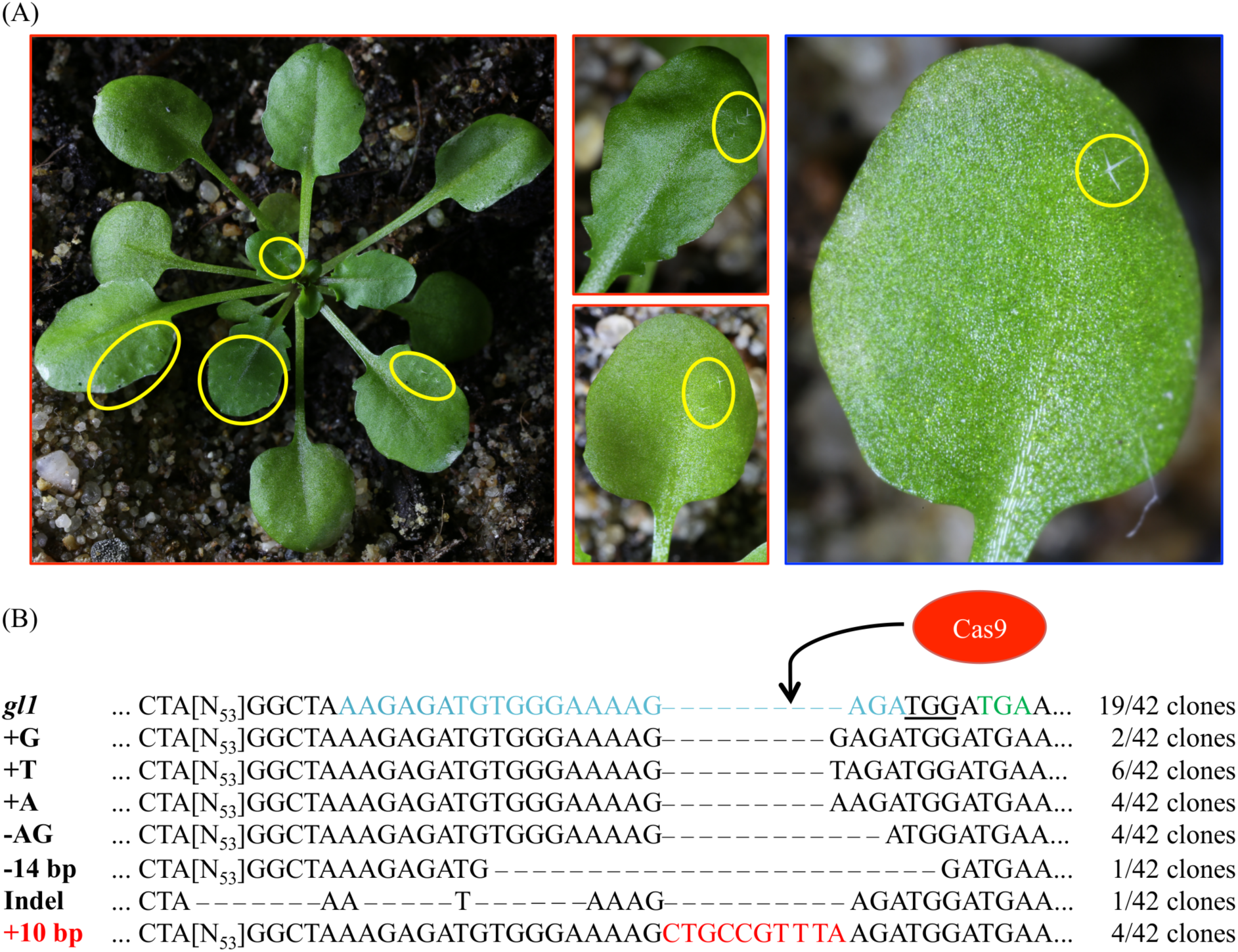
T2 generation plants show spots of trichomes. (A) Representative images of T2 plants transformed with pIPGT-Nuc (red frame) or pVIR-Nuc (blue frame) showing spots of trichomes (yellow circles) ranging from one single trichome to fully covered leaves. (B) The *gl1* gene was amplified from leaves with trichomes, subcloned and sequenced. Sequence analysis revealed clones with various indel mutations next to clones containing the desired 10 bp insertion (red letters). PAM sequence is underlined, STOP codon in *gl1* line is marked in green letters, Cas9 target site is marked in blue letters.

We isolated gDNA from leaf trichome spots of three IPGT plants (the single trichome on the pVIR-Nuc plant provided not enough plant material for gDNA isolation), amplified the *gl1* gene, subcloned the PCR product and sequenced 42 clones. While most clones still contained the *gl1* sequence with the 10 bp deletion, we detected various indel mutations but also the integration of the codon-modified 10 bp HT, which restored the wildtype amino acid sequence of the *GL1* gene (Fig. 3B). This chimerism on genotype level was in line with the highly chimeric phenotype of the leaves.

### Fully regenerated trichomes were detected in plants of the T3 generation

We harvested seeds from the chimeric T2 plants and analyzed ~ 200 progeny T3 plants of each T2 plant. In most cases, the progeny plants were glabrous or just contained single trichome spots. However, we detected three plants (frequency: 1.5 %) in the progeny of the highly chimeric IPGT plant (Fig. 3A, left) with fully regenerated trichomes (Fig. 4A). We amplified the *gl1* gene from those plants and sequenced the PCR product. Two of the three plants showed clear peaks corresponding to the 10 bp insertion but also peaks corresponding to small indel mutations (Fig. 4B, Fig. S3), indicating that only one of the two alleles was repaired by HR. In the third plant, we detected the expected 10 bp insertion not in the sequencing histogram but in 5 out of 15 sequenced clones of subcloned PCR product (Fig. S3D).

**Figure 4:**
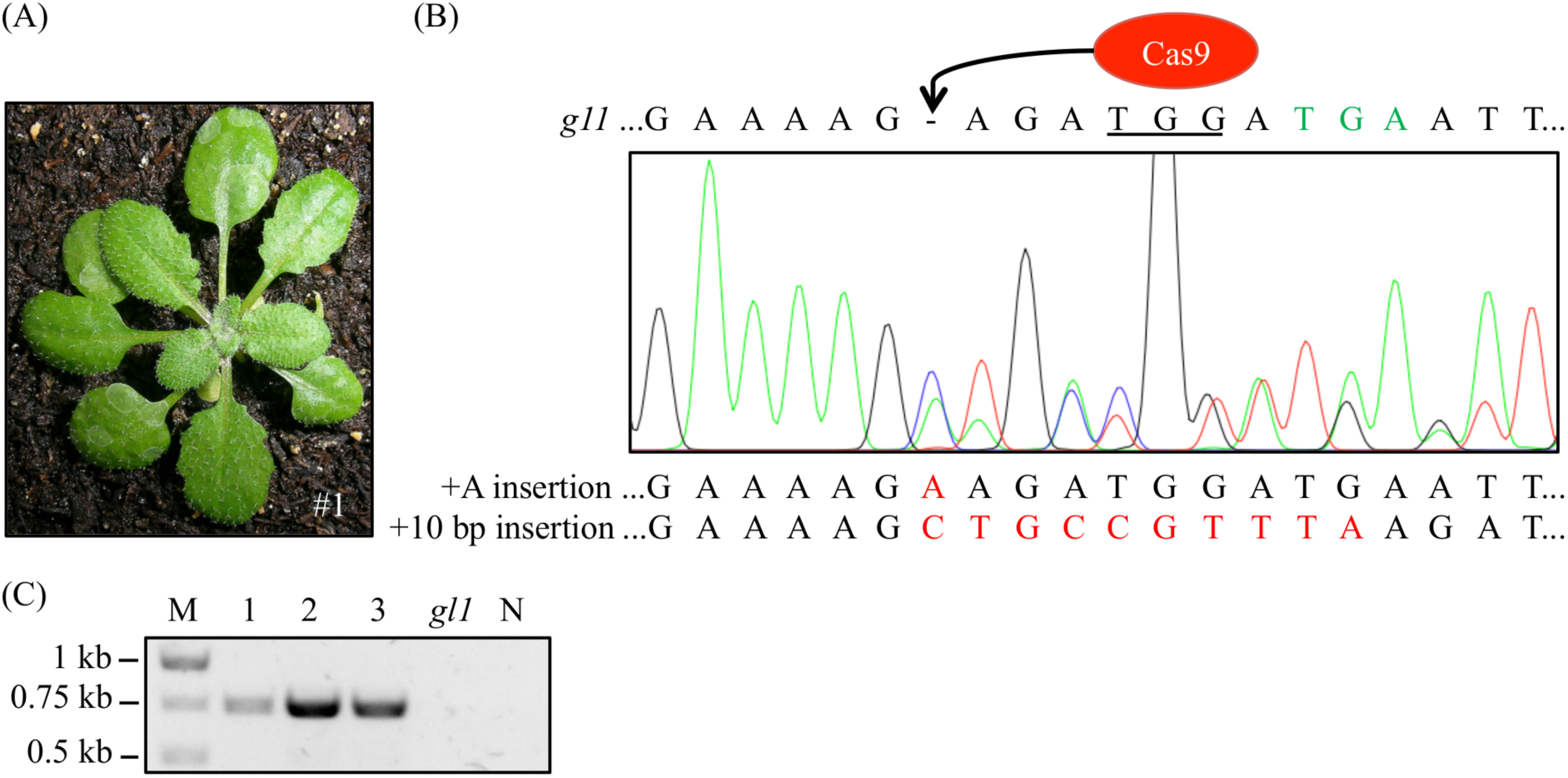
Wildtype-like trichome patterning was detected in three T3 generation plants. (A) Exemplary picture of one of the three T3 plants (#1) with full trichome covering. (B) We verified the repair of the *gl1* gene by amplifying the *gl1* gene and sequencing the PCR amplicon. Double peaks appeared at the site of Cas9 cleavage hinting at a chimeric or biallelic mutation, the peaks corresponded to an insertion of the ten bps (red) from the HT and to an adenine insertion. Sequence of the dysfunctional *gl1* gene is given on top of the sequencing histogram for comparison. Peak colors: Adenine = green, cytosine = blue, guanine = black, thymine = red; PAM sequence underlined, premature STOP codon marked in green letters. (C) Presence of the *Cas9* gene in the three non-glabrous plants was detected by PCR using *Cas9*-specific primers. All three plants revealed PCR amplicons at the expected size of 749 bp. In contrast, DNA from the *gl1* background line did not yield a PCR Cas9 signal. Image of agarose gel was color inverted for better visibility. M = Marker, N = water control.

### The repaired *GL1* allele segregates in a mendelian way

In genome edited plants, while the editing nuclease (Cas9) is active, it is hard to distinguish between somatic and heritable mutations. The three plants that fully recovered trichome production were still carrying the *Cas9* gene (Fig. 4C), which was likely still active. Thus, we confirmed in those plants that the 10 bp insertion is indeed heritable and not somatic by Mendelian segregation analysis. We analyzed the offspring of the three T3 plants displaying trichomes and counted the amounts of trichome-bearing and glabrous plants in the T4 generation. If one allele was indeed repaired by HR in the whole plant and the other one not, we would expect a Mendelian distribution of the trichome phenotype with a ratio of non-glabrous:glabrous plants of 3:1. Indeed, the progeny of all three plants did not deviate significantly from the expected 3:1 ratio as calculated by chi-squared test (χ^2^; P > 0.5, Tab. 2). Sequencing of 125 progeny plants from all three T3 plants with or without trichomes revealed that the trichome phenotype was always associated with at least one repaired *GL1* allele, while the glabrous plants mostly contained various small indel mutations (Fig. S3). Among the 125 sequenced plants, we detected 30 plants that showed a homozygous insertion of the 10 nucleotides according to the sequencing histogram (Fig. 5). The small indel mutations found in glabrous plants or in heterozygous non-glabrous plants were in many cases not consistent with the indel alleles found in the progenitor plants, indicating that only the repaired *GL1* allele was already fixed in the T3 generation and that the other sequenced allele in the T3 plants was caused by somatic mutations in the leaf used for gDNA extraction. This putative chimerism in the T3 generation plants corresponds to the presence of more than two sequence peaks in some of the sequencing histograms of the T3 parental plants (Fig. S3).

**Table 2:** Segregation pattern of trichome phenotype in T4 generation plants transformed with pIPGT-Nuc. Progenitor T3 plant number according to Fig. 4 and Fig. S3.

| Progenitor T3 plant | Number of plants with trichomes | Number of plants without trichomes | Ratio of plants with/without trichomes | $\chi^2$ value | P value |
| --- | --- | --- | --- | --- | --- |
| #1 | 164 | 55 | 2.98/1 | 0.0015 | > 0.9 |
| #2 | 150 | 52 | 2.88/1 | 0.0594 | > 0.5 |
| #3 | 210 | 75 | 2.80/1 | 0.2632 | > 0.5 |

**Figure 5.**
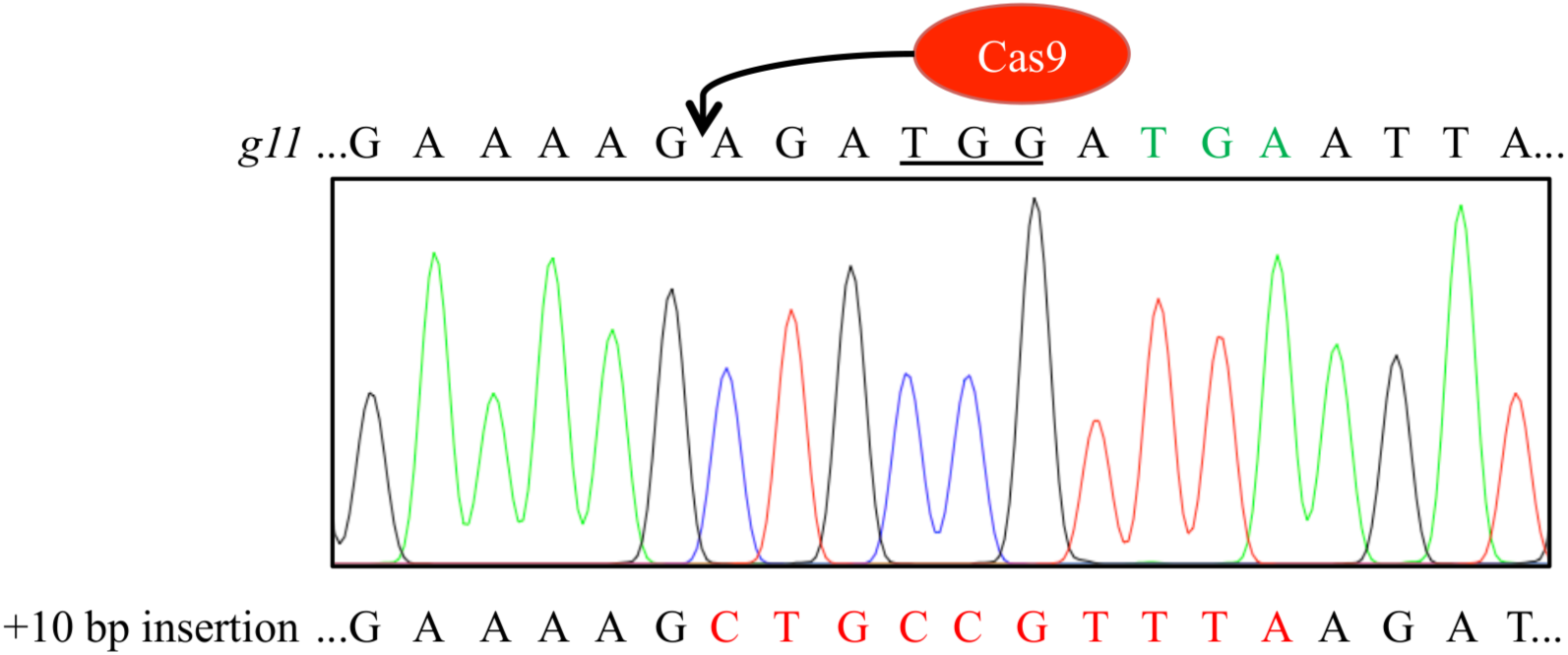
Sequencing histogram of T4 generation plant reveals homozygous integration of the 10 nucleotides thereby restoring the *GL1* reading frame. Sequence of the dysfunctional *gl1* gene is given on top of the sequencing histogram for comparison. Peak colors: Adenine = green, cytosine = blue, guanine = black, thymine = red; PAM sequence underlined, premature STOP codon marked in green letters.

In summary, we screened between 2,200 to 2,500 plants containing the *gl1* repair constructs pIPGT-Nuc, pVIR-Nuc, and pVIR-Nick, respectively, over three generations. Using the viral replicon approach, we could not regenerate non-glabrous plants. Using the IPGT approach, we were able to detect three stable GT events in the T3 generation with one functional *GL1* allele.

## DISCUSSION

### Trichome formation is a marker for mutagenic efficiency

The Cas9 system has become a valuable and efficient tool for genome modifications in plants (Arora and Narula, 2017). However, while random mutagenesis in plants by error-prone DSB repair through the NHEJ-pathway has been successfully applied in many studies (Liu et al., 2017), introduction of targeted mutations and gene knock-ins by GT remains challenging.

To explore the efficiency of two systems that have been described to induce HR in plants, we used a Cas9-generated trichomeless *Arabidopsis* mutant (Hahn et al., 2017). This mutant contains a 10 bp deletion (Fig S1) in the *GL1* gene, which leads to a premature STOP codon. The trichome phenotype (Fig. 3) allowed easy selection of transformed plants, which were likely to produce completely repaired offspring (Fig. 4).

An important aspect of our marker system is that DSB repair has to occur via HR and not via NHEJ to induce trichome formation. Small indels of 1–2 nucleotides generated by error-prone NHEJ might restore the open reading frame (ORF) of the gene, but at least three amino acid residues are still missing in the MYB-binding domain of the GL1 protein, which are critical for protein function (Hauser et al., 2001). As NHEJ mediated repair of DSB rarely results in insertions of more than one nucleotide (Fauser et al., 2014), it is highly unlikely that NHEJ mediated repair can restore trichome formation. Consequently, in a large-scale screening of glabrous and non-glabrous T4 plants (Fig. S4), we detected a clear correlation of trichome formation and repair via HR, while NHEJ mutations did not result in non-glabrous plants even though the ORF of the gene was partly restored in some cases. Previously used markers, such as ß-glucuronidase (GUS) or fluorescent proteins need first to be introduced into the plant and GT events can only be detected by staining or fluorescence microscopy. In contrast, our endogenous marker gene has two advantages: First, it allows to follow the mutagenesis progress by eye and second, trichome absence does not have pleiotropic effects on plant development under greenhouse conditions (Oppenheimer et al., 1991).

### Activation of the homology template by viral replicons did not result in gene targeting events

Baltes et al. (2014) developed a GT approach in tobacco that employs geminivirus-based replicons to active the HT (Fig. 2). In a first approach in *Arabidopsis*, they aimed at inserting a short DNA sequence into an endogenous gene by GT using a HT, which was activated by replication in recombinant bipartite cabbage leaf curl virus genomes. While GT events could be detected on somatic level by PCR, the authors did not report inheritance of the GT events to progeny plants (Baltes et al., 2014). In further experiments, several groups used deconstructed virus replicons of BeYDV (or other viruses) depleted of the coat and movement proteins. These can be stably transformed into plants and have an increased vector cargo capacity, allowing the use of larger HT. This approach allowed efficient GT in various crop plants, including tobacco, tomato, potato, rice and wheat (Baltes et al., 2014; Butler et al., 2016; Čermák et al., 2015; Gil-Humanes et al., 2017; Wang et al., 2017).

While the initial experiments with the full virus demonstrated that GT using viral replicons can be achieved in *Arabidopsis*, no further studies reported the use of deconstructed replicons in *Arabidopsis*. We were also unable to detect GT events in form of trichome regeneration in 2,334 analyzed plants from three independent lines transformed with pVIR-Nuc except a single trichome (Tab. 1, Fig. 3). One possible explanation could be that the sgRNA that we used did not allow efficient DSB induction in the target locus. Since we generated successful GT events with the IPGT approach, using the same sgRNA, which requires efficient Cas9 cutting for HT activation, we do not consider this as likely. As we detected high frequencies of NHEJ mutations next to GT events while sequencing T4 generation plants (Fig. S4), we conclude that our sgRNA is highly active. It could be possible that replicon formation is not efficient in *Arabidopsis* in contrast to other plant species. However, BeYDV infects *Arabidopsis* (Liu et al., 1997). Thus, replicon formation should be possible. We furthermore detected efficient replicon formation on DNA level in T1 plants (Fig. S2). Another option could be an *Arabidopsis*-specific discrimination of the repair machinery for circular HT as these do not occur naturally in the plant nucleus.

Differences between plant species with handling of foreign DNA molecules have already been observed in *Arabidopsis* and tobacco (Orel and Puchta, 2003). It has been suggested that the stimulatory effect of viral replicons on GT is not only caused by the amount of HT but also by the impact of RepA on the plant cell cycle. RepA stimulates entry of cells into the S-phase by interaction with plant proteins that control cell cycle so that the virus can use the necessary factors for replicon replication that the cell produces for DNA synthesis (Baltes et al., 2014). Cells in S-Phase show an increased HR efficiency in eukaryotes (Saleh-Gohari and Helleday, 2004; Mathiasen and Lisby, 2014). Possibly, this effect is less pronounced in *Arabidopsis* than in other plants.

In contrast to most of the studies using viral replicons, we did not include the SSN in the replicating unit. Baltes and colleagues (2014) reported comparable GT efficiencies if the ZFN was part of the replicating unit or not, and similar results were found in a Cas9-based approach in tomato (Čermák et al., 2015). However, it might be possible that a high amount of SSN is more critical in *Arabidopsis.*

Donor architecture might also account for variable efficiencies in GT. We used HA with a length of app. 800 bp, which is in the range of the most successful GT studies so far (Fauser et al., 2012; Baltes et al., 2014; Schiml et al., 2014; Čermák et al., 2015; Butler et al., 2016; Gil-Humanes et al., 2017; Wang et al., 2017). Additionally, we used the viral components described in the original report of Baltes et al. (2014) and just integrated our HT. Even though we used a different vector backbone (Fauser et al., 2014), the replicons should therefore be comparable to the ones described in successful GT studies by Baltes et al. (2014) and Čermák et al. (2015).

While we could not detect considerable trichome formation in our experiments, it has to be taken into account that GT events might occur within cells, which are not determined to develop into trichomes. That is, using trichome formation as readout for mutagenic efficiencies, we actually underestimate the amount of successful GT events on the somatic cell level. PCR-based methods, as used for GT detection in *Arabidopsis* using cabbage leaf curl virus genomes (Baltes et al., 2014) will therefore allow detection of more GT events. In other experiments of our group with a different target gene (unpublished data), we actually could detect GT events using PCR-based approaches on somatic level at low frequencies but we were also unable to regenerate inheritable events Taken together, these data indicate that viral replicons allow GT only at low frequencies in *Arabidopsis,* which contrasts with the considerable success in many crop species. Further experiments are needed to unravel the reasons for these differences so that this promising technique might also be applicable in this important model organism.

### Cas9 nickase is not improving HR events in combination with viral replicons

We tested whether Cas9 nickase can induce GT at higher efficiencies than Cas9 nuclease as proposed by Fauser et al. (2014). We only applied the Cas9 nickase in combination with the viral replicon approach, since IPGT needs a nuclease to release the HR template. We did not detect GT events in form of trichome formation on any plant (Tab. 1). However, when we combined the viral replicon approach with a Cas9 nuclease, we also detected only a single putative GT event in ~2,300 plants screened (Tab. 1, Fig. 3). Therefore, it is likely that the viral replicon approach does not trigger GT in *Arabidopsis,* independent from its Cas9 variant.

However, besides the initial report of Fauser et al. (2014), reports on GT using nickases in plants are scarce. For their GT experiments, Fauser and colleagues used a reporter construct (Orel et al., 2003) containing a disrupted *GUS* ORF as well as a homologous sequence, which can function as HT for repair of the *GUS* ORF upon induction of a DSB by Cas9. Up to 4 times more HR events were detected compared to experiments with Cas9 nuclease. Importantly, only somatic mutations were quantified and information on the regeneration of stably mutated plants was not provided. It has been observed earlier, that GT efficiencies increase if target locus and donor construct are located on the same chromosome (Fauser et al., 2012). In the GUS experiments described above, target locus and HT are directly adjacent, which possibly explains the successful HR events. In our case, the HT integrated randomly into the genome and even after replicon formation, the replicons still had to “find” the nick site to be available for HR. Supporting this hypothesis, other studies detected lower GT efficiency when using nickases compared to nucleases (Ran et al., 2013; Čermák et al., 2017). In a recent study, Cas9 nickase was used in combination with viral replicons to repair a dysfunctional *GUS* gene in tobacco. Using two sgRNAs to create adjacent nicks, a 2-fold increase in GT was observed (Čermák et al., 2017). However, paired nicks on different DNA strands in close proximity are considered to induce a DSB with single strand overhangs (Schiml et al., 2014). Thus, it is questionable if nick HR repair pathways were indeed responsible for the successful GT in that study. In human cells, DNA nick repair via HR occurs 8 times more frequently on the transcribed (non-coding) DNA strand, probably due to stimulation by the transcription machinery (Davis and Maizels, 2014). The Cas9 nickase used in our study only contains a functional HNH-cleavage domain so that the nick is created on the non-coding strand. However, whether the strand preference in plants is comparable to that in human cells remains to be elucidated and might provide a possibility to enhance HR frequencies.

In general, HR repair mechanisms on DNA nicks in plants are not well studied. In humans, two competing HR mechanisms are proposed. The canonical HR pathway resembles the synthesis-dependent strand annealing repair pathway for DSB, while the alternative one preferentially employs single stranded oligonucleotides or nicked dsDNA HT for repair (Davis and Maizels, 2014). The alternative HR pathway shares features with another HR pathway, the single strand annealing pathway, which is induced by micro-homologies at the break ends. As HR via the single strand annealing pathway seems to be more efficient in plants than the synthesis-dependent strand annealing repair pathway (Orel et al., 2003; Fauser et al., 2014), it might be interesting to include nickase target site in the repair template to induce target locus repair via the alternative HR nick repair pathway. However, the detailed mechanisms underlying DNA nick repair in plants via HR needs to be elucidated to verify if the same mechanisms as in human cells apply. As the Cas9 nickase has already been shown to induce gene knock in in human embryonic stem cells using dsDNA as template (Rong et al., 2014), it remains an interesting option for future HR experiments *in planta.*

### The *in planta* gene targeting approach allows regeneration of stable gene targeting events in Arabidopsis

The IPGT approach, which excises the HT from the T-DNA backbone by a SSN (Fig. 2), has been used successfully in *Arabidopsis.* By the application of the I-*Sce*I meganuclease a *GUS* marker gene was restored (Fauser et al., 2012), and by the application of the Cas9 system a resistance cassette was integrated into the *ADH* locus (Schiml et al., 2014) and a *GFP* gene introduced into the *TFL1* locus (Zhao et al., 2016). In contrast to these previous studies, we did not exploit an artificial marker gene but restored the gene function of the endogenous *GL1* gene, using trichome formation as direct read-out for GT events.

We detected spots of trichomes ranging from single trichomes to fully covered leaves in T2 generation plants (Fig. 3, Tab. 1). Other studies reported similar chimeric GT spots using GUS staining (Fauser et al., 2012). The T2 plant, which showed the highest level of chimerism (Fig. 3) produced offspring plants with WT-like trichome patterning. We therefore conclude, that the trichome phenotype can be used to select for plants, in which GT efficiencies are enhanced and which are candidate plants for production of stably mutated progeny. This enhancement can have several reasons. The location of the T-DNA, which contains HR template and SSN gene, in the plant genome likely has influence on GT and close proximity of repair template and DSB locus seems favorable (Fauser et al., 2012). Similarly, we found differences in the frequency of chimeric T2 plants depending on the parental T1 line (Tab. 1). It is therefore highly recommended to start with as many independent transformation events as possible, in contrast to screening a lot of progeny plants from few primary transformants (Schiml et al., 2017).

T2 plants might be homozygous or heterozygous for the T-DNA insertion. It has been shown that the heterozygosity state of the HR template influences recombination frequencies (Puchta et al., 1995; Orel et al., 2003). Indeed, since all three plants, which completely restored trichome formation in the T3 generation (Fig. 4, Fig. S3), contained the Cas9-T-DNA cassette and also produced only T-DNA containing offspring, it is likely, that the parental T2 plant contained already two copies of the repair template. Crop improvement usually requires the creation of marker-free plants. Thus, more screening effort might be needed to detect T-DNA free GT plants.

It has been reported that one-sided GT events can occur during repair (Schiml et al., 2014), in which one side of the of the DSB is repaired by HR and the other strand is repaired using the stably integrated T-DNA as template. This can lead to integration of large parts of the HT, which finally results in a NHEJ-mediated ligation of the repaired strand and the loose end of the DSB. To detect such events, we had added single bp exchanges in both HA. Our sequencing results of our repaired *GL1* alleles did not contain these bp exchanges, which indicates that we achieved true GT events.

In previous studies the frequencies of GT events using IPGT in *Arabidopsis* ranged from 0.14–0.83 % (Fauser et al., 2012; Schiml et al., 2014; Zhao et al., 2016) which corresponds well to our GT frequency of 0.12 %. The single bp exchanges in the HA, which allowed us to detect true GT events, might explain the slightly lower efficiencies in our experimental approach. Additionally, efficiencies in Cas9 experiments are difficult to compare due to use of different sgRNAs (Peng et al., 2016).

### Further options to enhance GT

In this report, we identified three stable GT event out of ~2,500 *Arabidopsis* plants screened across three generations using an IPGT approach. In principle, a GT event at such frequency could be detected exclusively by PCR and without a phenotype based screening. However, we achieved such a high frequency because we were able to select parental lines prone to GT by phenotype. Therefore, it might still be desirable to increase GT efficiencies if no visual marker is used for GT detection. In general, it is advisable to test sgRNA efficiencies in protoplast systems before using them in stable plant transformations as efficient induction of DSB is a prerequisite for successful GT (Puchta et al., 1993).

Higher GT efficiencies of ~5.5 % have been achieved in *Arabidopsis* protoplasts using single stranded oligonucleotides as repair template (Sauer et al., 2016) to exchange specific nucleotides in a fluorescent protein to change its emission spectrum. However, single stranded oligonucleotides are not suitable for knock-in of larger DNA fragments or whole genes due to size limitations and are not transformed via Agrobacteria. For targeted exchange of a single bp, the use of base changer Cas9 provides an attractive alternative to HR-mediated repair processes. In this approach, an inactive Cas9 is fused to cytidine deaminase, which converts cytosines to uracil without DNA cutting (Komor et al., 2016; Zong et al., 2017). Recently, an adenine base editor Cas9 has also been described (Gaudelli et al., 2017). While mutation frequencies of up to 50% can be achieved using base changer Cas9, this approach cannot be applied for gene knock-ins.

In general, protoplast transformation seems to yield higher mutation frequencies compared to Agrobacterium-mediated transformation (Li et al., 2013; Sauer et al., 2016), since more DNA donor molecules are integrated in the plant cells. However, regeneration of *Arabidopsis* plants from protoplasts is challenging and is therefore not an attractive alternative to floral dipping. Focusing research on improved regeneration protocols is therefore desirable. Several groups tried to enhance HR efficiencies by disrupting either the NHEJ pathway (Maruyama et al., 2015; Zhu et al., 2015) or introducing HR stimulating enzymes from other organisms (Reiss et al., 2000; Shaked et al., 2005; Even-Faitelson et al., 2011). These approaches can destabilize the genome as HR frequencies are enhanced between all repetitive sequences in the genome (Puchta and Fauser, 2013).

In the IPGT approach, the HT is excised from the genome. If this happens at an early stage of plant development, the HT might not be passed on during cell division. This could reduce the efficiency of this approach. Using germline-specific promoters (Wang et al., 2015; Mao et al., 2016) for the SSN might increase chances of simultaneous cleavage of target locus and release of the HT specifically in the cells, where GT needs to occur to generate stable offspring.

A similar problem is target locus inactivation through error-prone NHEJ repair. Besides the before mentioned nickase approach, the Cas9 variant CpfI (CRISPR from *Prevotella* and *Francisella*) might provide an attractive alternative. CpfI generates DSB more distal to the PAM than Cas9 (Zetsche et al., 2015), thus making the target locus accessible to a second round of cleavage even after NHEJ mediated repair. An interesting approach would be to combine the viral replicon approach with the IPGT approach. Therefore, the HT would be flanked by SSN cutting site and the viral components. By this, large amounts of linear dsDNA molecules would be generated that can be used as repair templates. Since more target sites are present in this approach, we suggest including the gene for the SSN and the sgRNA within the replicon unit, so that sufficient amounts of all genome editing components are available.

## CONCLUSION

In this report, we applied the Cas9 system to induce GT events in *Arabidopsis.* The HT needed to be activated by an IPGT approach. We were able to stably introduce a short DNA sequence into an endogenous gene and the inserted sequence was transmitted through the germline into the next generation. The high GT efficiency allows to use GT approaches also in marker-free approaches in which GT events need to be detected by PCR. Thus, we conclude that Cas9 mediated GT in combination with IPGT is a powerful tool that has high potential to become routinely applied in plant research in the future. Trichome formation as marker for editing efficiency proved to be a reliable system to compare different GT approaches. We suggest that the *GL1* gene can be used for optimization of Cas9-based mutagenesis approaches in future experiments.

## AUTHOR CONTRIBUTIONS

FH, OM, ME, and AW designed the study. FH carried out all laboratory experiments. FH, OM, ME, and AW interpreted the data and wrote the manuscript.

## ABBREVIATIONS

BeYDV: *bean yellow dwarf virus*
bp: base pair(s)
Cas9 system: CRISPR/Cas9 system
DSB: double strand break
GT: gene targeting
GUS: ß-glucuronidase
HA: homology arm(s)
HR: homologous recombination
HT: homology template
IPGT: *in planta* gene targeting
LIR: long intergenic region
NHEJ: non-homologous end joining
ORF: open reading frame
PAM: protospacer adjacent motif
sgRNA: single guide RNA
SIR: short intergenic region
SSN: sequence specific nuclease(s)
TALEN: Transcription Activator-like Effector Nuclease
ZFN: Zinc finger nuclease(s)

## ACKNOWLEDGMENTS

We thank Prof. Dr. Holger Puchta and Felix Wolters for providing us with the vectors pDE-Cas9, pDE-Cas9-D10A and pEN-C1.1, and for valuable feedback. We acknowledge Prof. Dr. Peter Westhoff and Dr. Shizue Matsubara for helpful discussions. We thank Steffen Köhler for photography support. We thank the gardeners for plant care and the student helper team for their support.

## CONFLICT OF INTEREREST STATEMENT

The authors declare that the research was conducted in the absence of any commercial or financial relationships that could be construed as a potential conflict of interest.

